# FGF9, a potent mitogen, is a new ligand for integrin αvβ3, and the FGF9 mutant defective in integrin binding acts as an antagonist

**DOI:** 10.1101/2023.12.01.569657

**Authors:** Chih-Chieh Chang, Yoko K Takada, Chao-Wen Cheng, Yukina Maekawa, Seiji Mori, Yoshikazu Takada

## Abstract

FGF9 is a potent mitogen and survival factor, but FGF9 protein level is generally low and restricted to a few adult organs. Aberrant expression of FGF9 usually results in cancer. However, the mechanism of FGF9 action has not been fully established. Previous studies showed that FGF1 and FGF2 directly bind to integrin αvβ3 and this interaction is critical for signaling functions (FGF-integrin crosstalk). FGF1 and FGF2 mutants defective in integrin binding were defective in signaling, whereas the mutants still bound to FGFR, and suppressed angiogenesis and tumor growth, indicating that they act as antagonists. We hypothesize that FGF9 requires direct integrin binding for signaling. Here we show that docking simulation of interaction between FGF9 and αvβ3 predicted that FGF9 binds to the classical ligand-binding site of αvβ3. We showed that FGF9 actually bound to integrin αvβ3, and generated an FGF9 mutants in the predicted integrin-binding interface. An FGF9 mutant (R108E) was defective in integrin binding, activating FRS2α and ERK1/2, inducing DNA synthesis, cancer cell migration, and invasion in vitro. R108E suppressed DNA synthesis induced by WT FGF9 and suppressed DNA synthesis and activation of FRS2α and ERK1/2 induced by WT FGF9 (dominant-negative effect). These findings indicate that FGF9 requires direct integrin binding for signaling and that R108E has potential as an antagonist to FGF9 signaling.

## Introduction

Integrins are a superfamily of cell adhesion receptors and recognize extracellular matrix (ECM) ligands, cell surface ligands, and small soluble ligands (e.g., growth factors) [1]. It has been established that integrins are critically involved in growth factor signaling through integrin-growth factor crosstalk [2]. The first indication of the role of integrins in growth factor signaling is that antagonists to integrin αvβ3 suppressed FGF2-induced angiogenesis [3]. The specifics of this crosstalk are, however, unclear. Currently popular models of integrin and growth factor crosstalk suggest that integrins promote growth factor signals through interaction of integrins with the extracellular matrix [4].

The fibroblast growth factor (FGF) family is comprised of 22 members that can be divided into seven subfamilies in mammals [5]. FGF9 (FGF9 subfamily) is a secretory protein that was first isolated from human glioma cells [6]. Four isoforms of FGF receptors mediate FGF9’s biological effects [5]. FGF9 binding to its receptor causes receptor dimerization and activates different signal transduction cascades [7, 8]. FGF9 is involved in a variety of complex responses in human or animal development. Mice lacking *Fgf9* have hypoplastic lungs [9], sex reversal [10], impaired germ cells [11], impaired skeletal growth [12], defective cardiomyocyte growth, and impaired inner ear development [13, 14].

Although FGF9 messenger RNA (mRNA) is ubiquitously expressed in embryos, FGF9 protein expression is generally low and restricted to a few adult organs [15]. Aberrant activation of FGF9/FGFR signaling is associated with cancers [16-19]. FGF9 has oncogenic activity and is involved in the progression of cancers in lung [15], stomach [20], colon, testis [16] and ovary [21]. FGF9 promotes epithelium and mesenchyme proliferation in lung [22]. Also, FGF9 enhances cell proliferation and invasive ability of prostate cancer cells [23] and ovarian cancer [21]. Overexpression of FGF9 can promote tumor growth and liver metastasis of mouse Lewis lung carcinoma via EMT induction [24]. In addition, elevated FGF9 expression is associated with poor prognosis in non-small cell lung cancer (NSCLC) [25].

Previous studies showed that FGF1 signaling requires direct integrin binding and subsequent integrin-FGF1-FGFR1 ternary complex formation is required for signaling functions and an FGF1 mutant defective in integrin binding (R50E) was defective in signaling functions [26] and suppressed FGF1 signaling induced by WT FGF1 act as an antagonist of FGFR [27]. FGF1 mutants defective in integrin binding strongly blocked angiogenesis in vitro and tumor growth in vivo [27, 28]. Similar results were obtained in FGF2 [28]. It is unclear if FGF9 requires crosstalk with integrins for signaling. We hypothesized that FGF9 directly binds to integrins and that FGF9 mutants defective in integrin binding acts as antagonists for FGF9 signaling.

In the present study, we showed that FGF9 directly binds to integrin αvβ3. Docking simulation predicted that FGF9 bound to the classical ligand-binding site of integrin αvβ3. We introduced point mutations in the predicted integrin binding interface of FGF9. An FGF9 mutant R108E was defective in integrin binding and in signaling functions. We found that the R108E suppressed signaling induced by WT FGF9, indicating that R108E is a dominant-negative antagonist of FGF9. Thus, we propose that R108 has therapeutic potential and the integrin-FGF9 interaction is a novel therapeutic target.

## Results

### FGF9 directly binds to integrin αvβ3

We found that soluble integrin αvβ3 (extracellular domains) bound to immobilized FGF9 in a dose-dependent manner in ELISA-type binding assays in 1 mM Mn^2+^ (Fig. 1a), as predicted by docking simulation. Cyclic RGDfV, an inhibitor specific to αvβ3, suppressed the binding of αvβ3 to FGF9, indicating that αvβ3 specifically bound to FGF9 (Fig. 1b). The binding of FGF9 to integrin αvβ3 required Mn^2+^, but Ca^2+^, Mg^2+^ or EDTA (all 1 mM) did not support FGF9 binding tp integrin αvβ3 (Fig 1c). Also, heat treatment reduced FGF9 binding to αvβ3, indicating that FGF9 should be properly folded. These findings indicate that FGF9 is a ligand for integrin αvβ3.

**Fig. 1.**
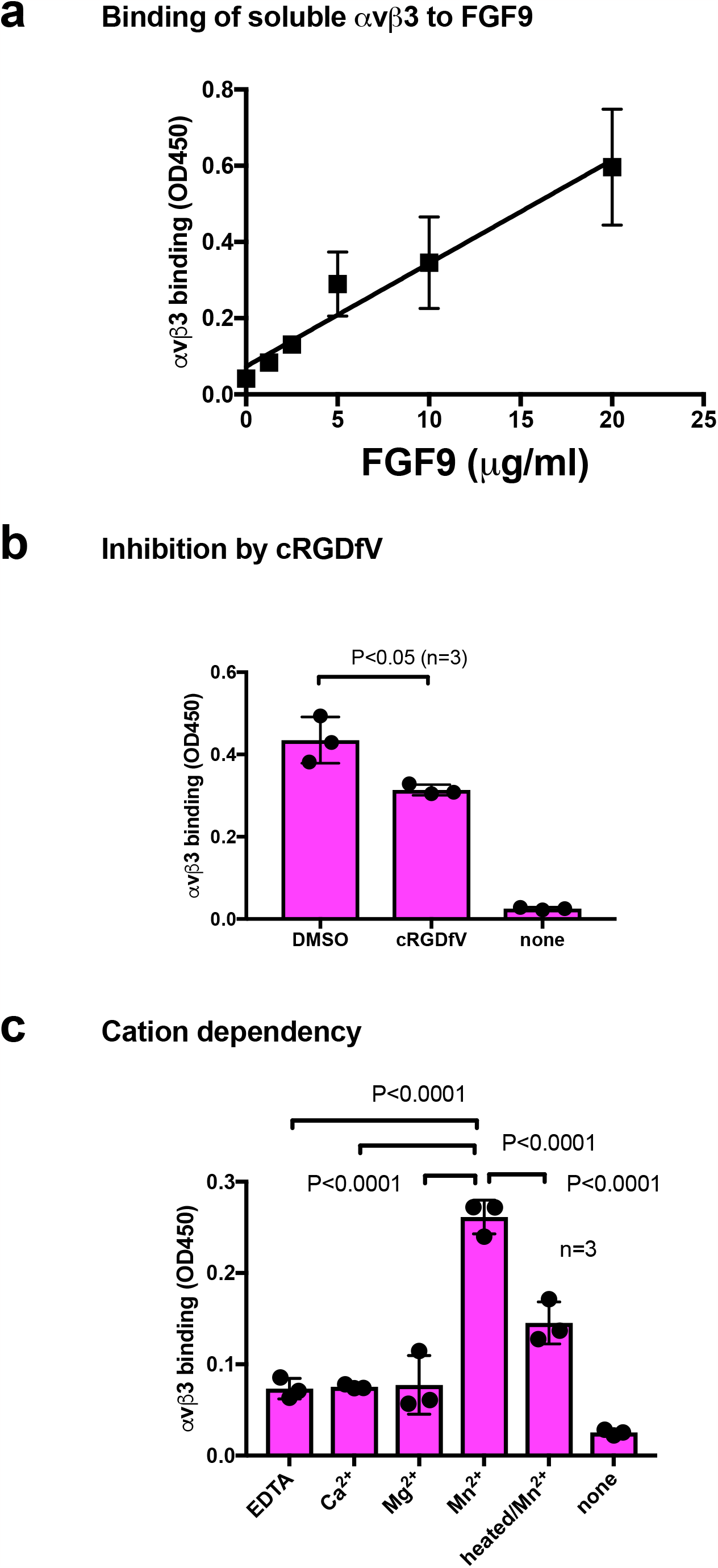
Binding of soluble integrin αvβ3 to FGF9. a. Binding of FGF9 mutant to αvβ3 in ELISA binding assay. Wells of 96-well microtiter plate were coated with FGF9 and remaining protein binding sites were blocked with BSA. Soluble αvβ3 (1 μg/ml) was added to wells and incubated for 1 h in Tyrode-HEPES buffer containing 1 mM Mn^2+^. Bound αvβ3 was quantified using anti-β3 and HRP-conjugated anti-mouse IgG. b. Binding of soluble αvβ3 in the presence of cyclic RGDfV, a specific inhibitor for αvβ3. Data are expressed as means ± S.D of triplicate experiments. c. The effectof cations and heat treatment on the binding of soluble integrin αvβ3 to FGF9. Cations (1 mM) was included in Tyrode-HEPES buffer. Data are expressed as means ± S.D of triplicate experiments.

To study how FGF9 binds to integrin αvβ3, we performed docking simulation of interaction between αvβ3 (PDB code 1L5G) and FGF9 (PDB code 1IHK) using AutoDock 3. The simulation predicted that FGF9 binds to the classical ligand binding site of αvβ3 at high affinity (docking energy -23 Kcal/mol, the first cluster) (Fig. 2a). When the 3D structure of the FGF9-FGFR1 complex (5W59.pdb) was superposed with the αvβ3-FGF9 docking model, there was little or no steric hindrance (Fig. 2b). This predicts that the FGFR1-binding site and the integrin-binding site are distinct. Amino acid residues that are predicted to be involved in FGF9-integrin αvβ3 interaction are shown in Table 1.

**Table 1.** Amino acid residues that are predicted to be involved in integrin αvβ3 and FGF9 interaction (site 1).

| $\alpha\text{v}$ | $\beta 3$ | FGF9 |
| --- | --- | --- |
| Asp146, Asp148, Ala149, Asp150, Tyr178, Trp179, Gln182, Asn205, Asn206, Gln207, Leu208, Ala209, Arg211, Thr212, Ala213, Gln214, Ala215, Phe217, Asp218, Asn260, | Ser121, Tyr122, Ser123, Met124, Lys125, Asp126, Asp127, Leu128, Trp129, Tyr166, Cys177, Tyr178, Asp179, Met180, Lys181, Yhr182, Arg214, Asn215, Arg216, Asp217, Ala218, Pro219, Glu220, Asp251, Thr311, Glu312, Asn313, Leu333, Ser334, Met335, Asp336, Ser337 | Thr52, Asp53, Asp55, His56, Leu57, Pro78, Asn79, Gly80, Thr81, **Arg108**, Val110, Asp111, Ser112, Gly113, Leu114, Tyr115, Asn119, Glu120, Tyr125, Gly126, Ser127, Glu128, **Lys129**, Leu130, Thr131, Gln132, Glu133, Cys134, Asn151, Leu152, **Lys154**, Asp203, Leu204, Leu205, Ser206, Glu207, Ser208 |

Amino acid residues within 0.6 nm between HBD and $\alpha\beta 3$ were selected using Pdb Viewer (version 4.1). Amino acid residues in HBD that are selected for mutagenesis are shown in bold.

**Fig. 2.**
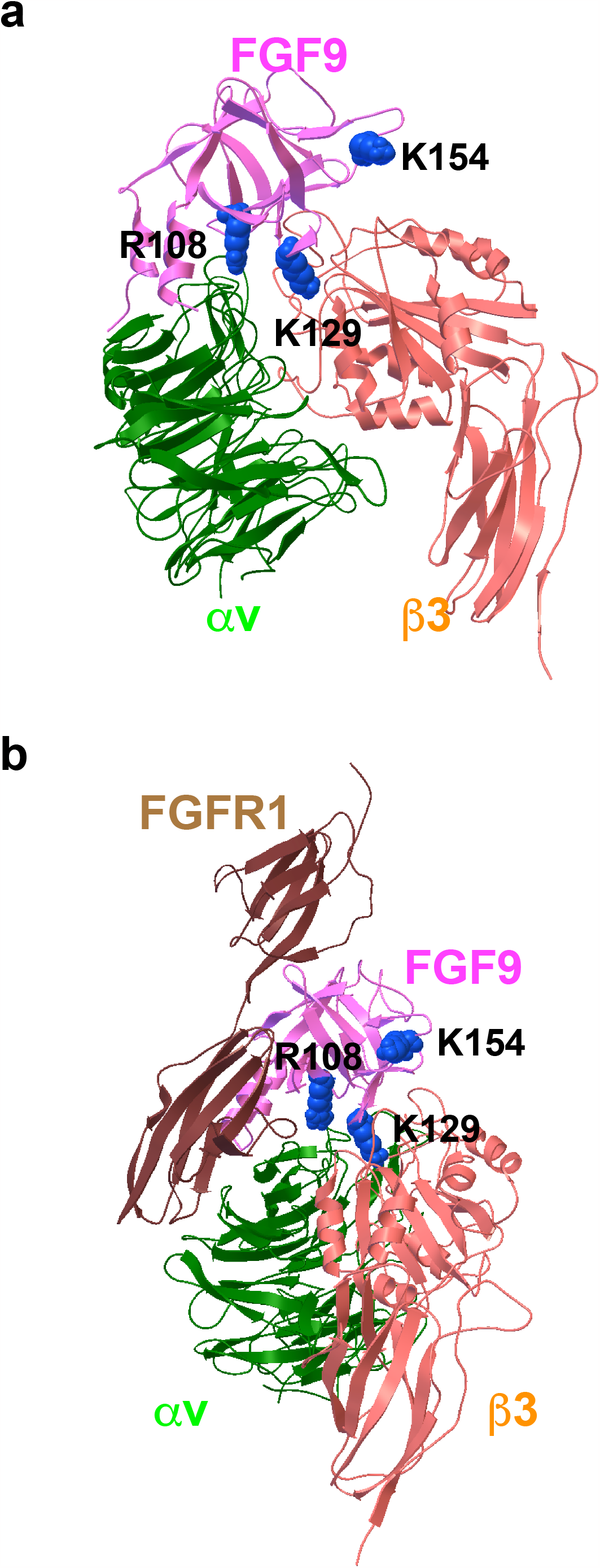
Docking simulation of FGF9-integrin interaction. Docking simulation of interaction between FGF9 and integrin αvβ3 predicted that FGF9 binds to αvβ3 at high affinity (docking energy -23 Kcal/mol). a. The FGF9-αvβ3 docking model. b. FGF9-FGFR1 complex was superposed to the FGF9-αvβ3 docking model. The simulation predicted that FGF9 binds to the classical ligand-binding site of αvβ3.

### Generation of Integrin-binding defective FGF9 mutants

We chose Arg108, Lys129 and Lys154 in the predicted integrin-binding interface of FGF9 for mutagenesis studies (Fig. 1a, Table 1). The Arg108 to Glu (the R108E mutation) and the K154E mutations and much less the K129E mutation reduced the binding of soluble αvβ3 to FGF9 in ELISA-type binding assays (Fig. 3b). This result indicated that the R108E and K154E mutations effectively suppressed the binding of FGF9 to αvβ3. We studied if FGF9 mutations affect DNA synthesis in NIH3T3 cells. (Fig. 3c). R108E completely lost the ability to induce BrdU incorporation, but K129E and K154E mutants were partly defective, indicating that direct binding to integrin αvβ3 is critical for signaling functions of FGF9. We selected R108E for further analysis.

**Fig. 3.**
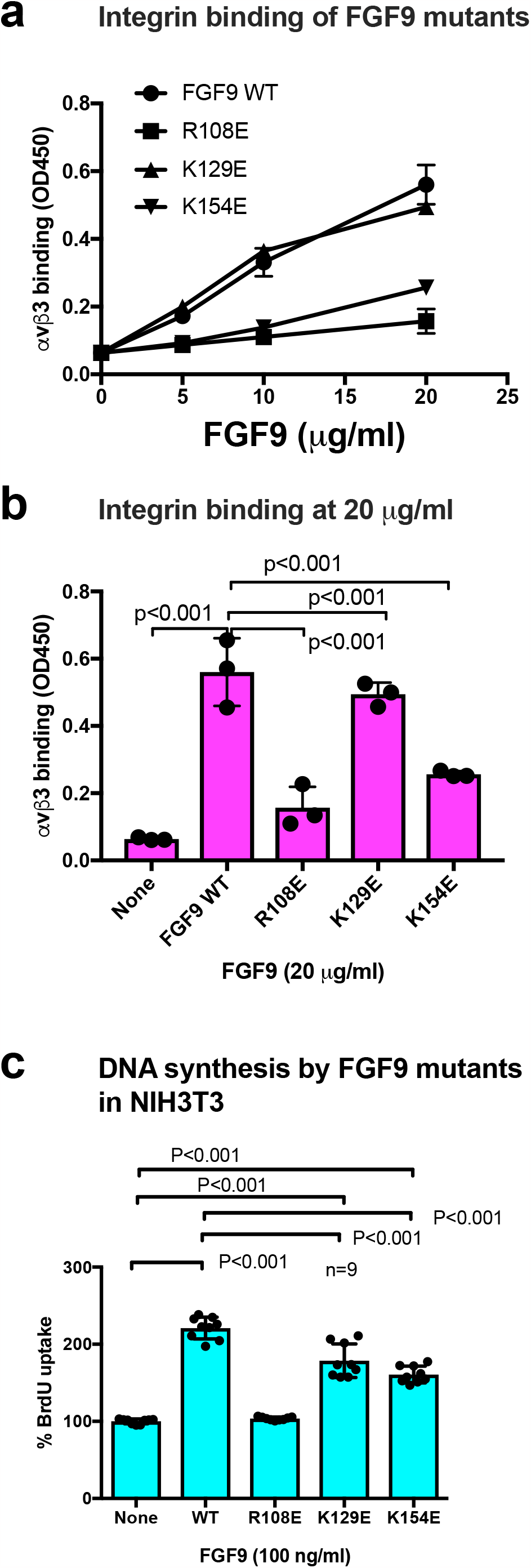
Identification of FGF9 mutants defective in integrin binding. a. FGF9 mutants are defective in integrin binding. We chose several amino acid residue of FGF9 for mutagenesis studies. Arg108, and Lys129, and Lys154 of FGF9 in the predicted integrin-binding site (Fig. 2) were mutated to Glutamate. The ability of FGF9 mutants (the R108E, K129E, and K154E mutants) to integrin was measured in binding assays using soluble αvβ3. Data are shown as means +/-SD of triplicate experiments. b. The binding of FGF9 mutants at 20 μg/ml is shown. Data are shown as means +/-SD of triplicate experiments. c. Effect of FGF9 mutations on DNA synthesis. NIH3T3 cells were starved for 24 hrs and stimulated with FGF9 (WT and mutants at 100 ng/ml) and BrdU was added to the medium for the last two hrs of the incubation.

### R108E is defective in inducing migration and invasion of colon cancer cells

DLD1 human colon cancer cells express FGFR2 and FGFR3, and Colon26 mouse colon cancer cells express FGFR3 (Fig. 4a). We studied the ability of WT FGF9 and R108E to induce pERK1/2. Fetal bovine serum (FBS, 10%) was used as a positive control. WT FGF9 induced ERK1/2 activation, but R108E was defective in inducing ERK1/2 activation (Fig. 4b). FGF9 is a potent chemoattractant. We found that R108E was defective in inducing cell migration (Figs. 4c and 4d). R108E was also defective in inducing invasion of DLD1 and Colon26 cells (Fig. 4e and 4f). These findings suggest R108E is defective in inducing migration and invasion in colon cancer cells.

**Fig. 4.**
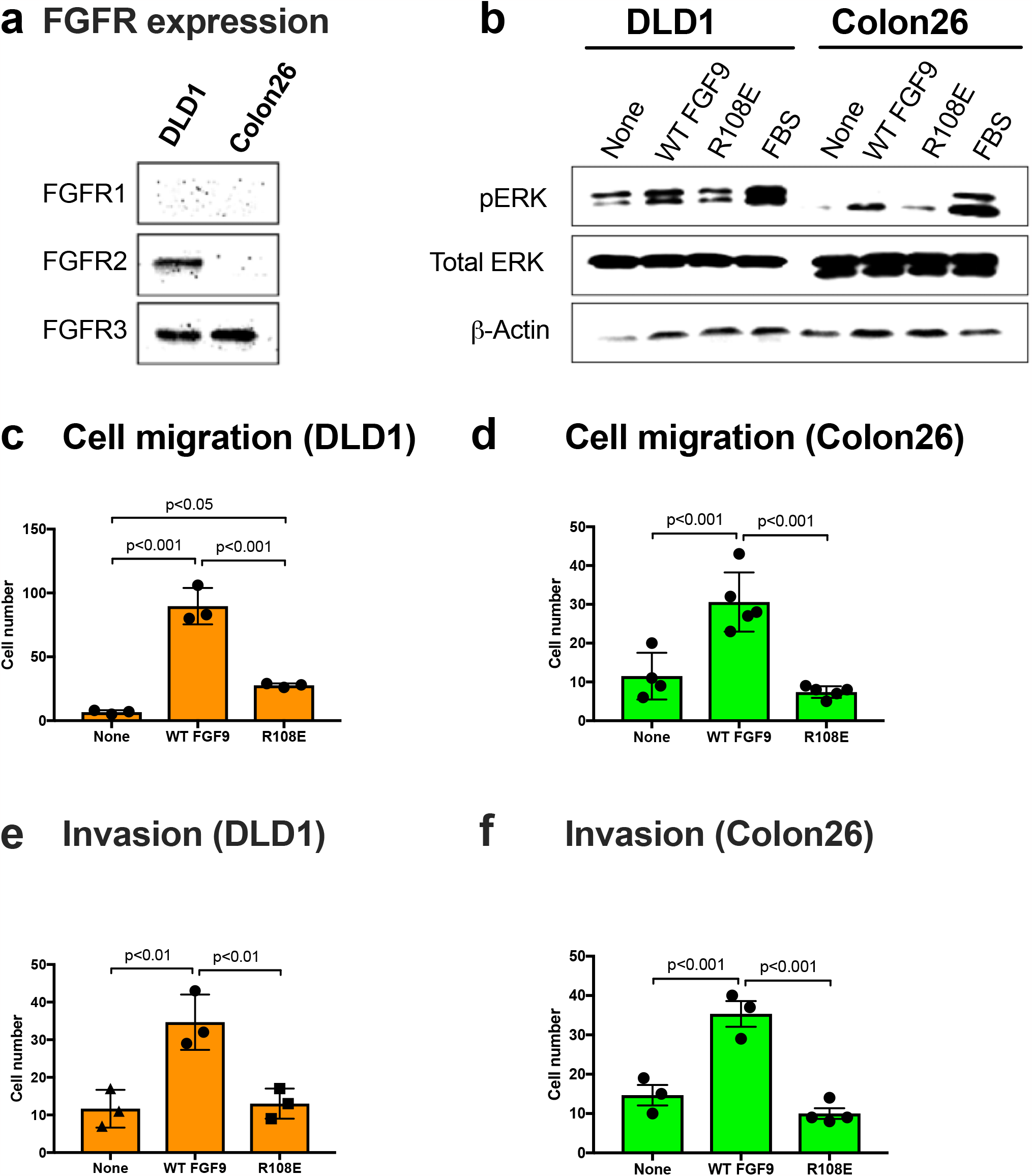
FGF9 mutant defective in integrin binding (R108E) is defective in inducing cell migration and invasion. a. Expression of FGFRs in DLD1 and Colon26 colon cancer cells. Cell lysates were analyzed by western blotting with antibodies specific to FGFR1, FGFR2, and FGFR3. b. FGF9-induced ERK1/2 activation in DLD1 and Colon26 colon cancer cells. DLD1 and Colon26 cells were serum starved for 24 hrs and stimulated with WT FGF9 (50 ng/ml) or R108E (50 ng/ml). Cell lysates were analyzed by western blotting. c. and d. Migration of DLD1 and Colon26 colon cancer cells to FGF9 was measured using Chemotaxicell chamber as described in the method section. Data are shown as means +/-SD of triplicate experiments. e.and f. Invasion of DLD1 and Colon26 colon cancer cells induced by FGF9 was measured using Chemotaxicell chamber coated with growth factor-depleted Matrigel as described in the method section. Data are shown as means +/-SD of triplicate experiments.

### R108E suppresses activation of FRS2 and ERK1/2 and DNA synthesis induced by WT FGF9 (dominant-negative effect) in NIH3T3 cells

If integrin binding to FGF9 and integrin-FGF9-FGFR ternary complex formation is required for FGF9 signaling, it is expected that FGF9 R108E mutant defective in integrin binding is expected to be dominant-negative, as in the case of FGF1 and FGF2 [27, 28]. We studied if R108E suppresses FRS2α and ERK1/2 phosphorylation induced by WT FGF9 in NIH3T3 cells. NIH3T3 cells were incubated with WT FGF9 and/or R108E for 30 min. We found that 20-fold excess R108E blocked FRS2α and ERK1/2 activation induced by WT FGF9 (Fig. 5a-c). We examined whether R108E suppresses DNA synthesis induced by WT FGF9 in NIH3T3 cells in BrdU incorporation assays. R108E did not induce DNA synthesis. We found that excess (20x) R108E suppressed DNA synthesis induced by WT FGF9 (Fig. 5d). These findings suggest that R108E is a dominant-negative mutant of FGF9.

**Fig. 5.**
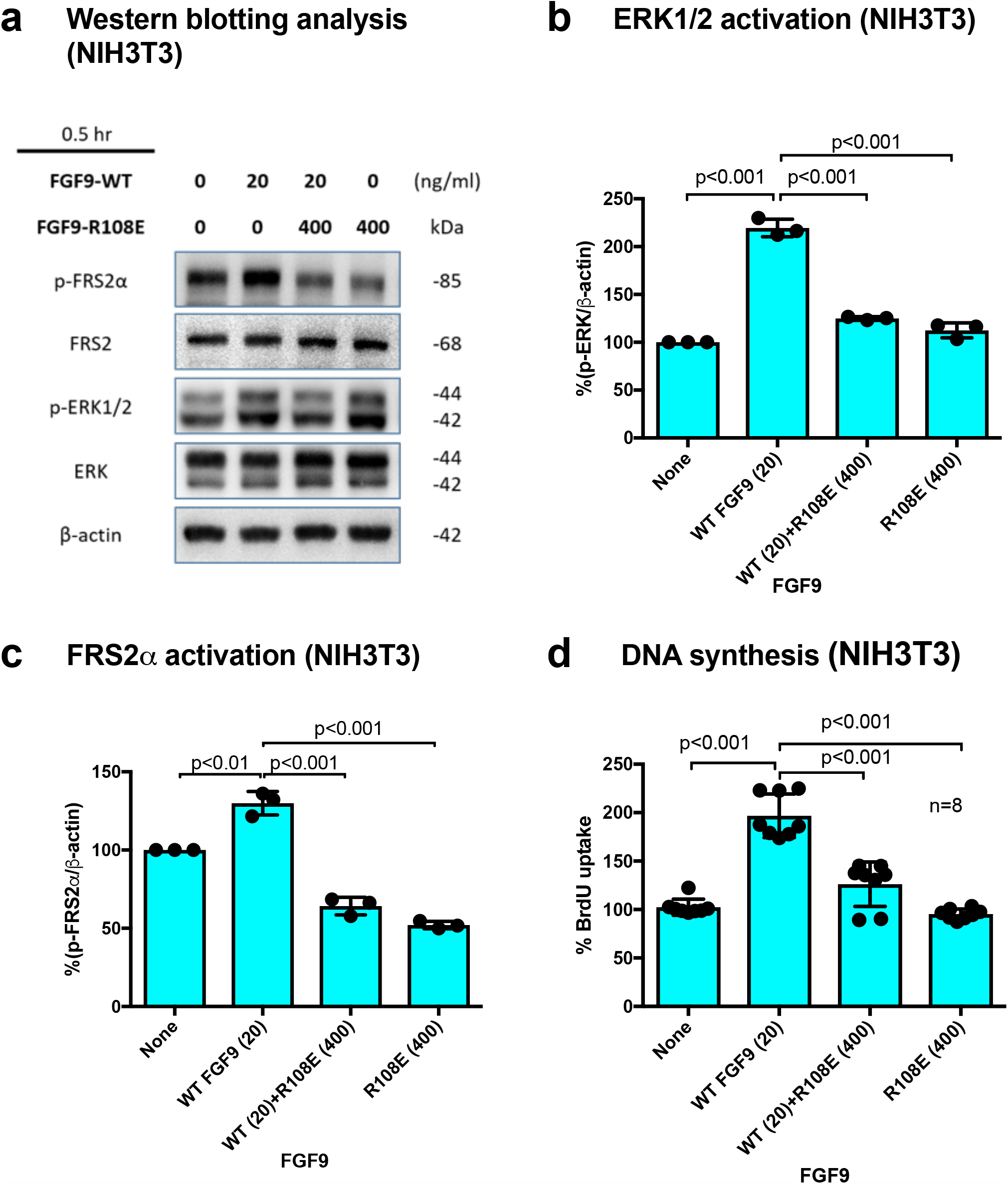
R108E suppresses FRS2 and ERK1/2 activated by WT FGF9 in NIH3T3 cells. a. NIH3T3 cells were starved for 24 hrs and stimulated with WT FGF9 (20 ng/ml) and/or R108E (400 ng/ml). Cell lysates were analyzed by Western blotting using anti-phospho-FRS2α, anti-FRS2α, anti-phospho-ERK1/2 and anti-total ERK1/2. β-actin was used as an internal control. Density of the bands were quantified using ImageJ software and phospho-ERK1/2/β-actin (b) or phospho-FRS2α/β-actin (c) was calculated. Data are shown as means +/-SD of triplicate experiments. d. NIH3T3 cells were starved for 24 hrs and stimulated with WT FGF9 (20 ng/ml) and/or R108E (400 ng/ml) and BrdU was added to the medium for the last two hrs of the incubation. Data are shown as means +/-SD of triplicate experiments.

## Discussion

### R108E (FGF9 antagonist) has potential as therapeutic in cancer

FGF9 has been implicated in the pathogenesis of cancer through its high affinity receptor FGFR3c. FGF9 is over-expressed in cancer-associated fibroblasts (CAFs) and mediates communication with cancer cells [29]. This indicates that FGF9 is a major therapeutic target in cancer, and antagonists to FGF9 have to be developed. In our previously studies, docking simulation predictes that FGF1 and FGF2 directly bind to integrins and we developed dominant-negative FGF1 and FGF2 mutants, which have therapeutic potential [26, 28]. The present study for the first time showed that FGF9 requires direct integrin binding for signaling functions. Docking simulation predicted that FGF9 binds to the classical ligand-binding site of αvβ3. We generated an FGF9 mutant defective in integrin binding (the R108E mutant). The position of R108E mutation cannot be deduced from comparison of primary structure of FGF9 and FGF1/FGF2, probably because FGF1, 2 and 9 interact with integrins differently.

The R108E mutant was defective in signaling functions, including DNA synthesis, activation of ERK1/2 and FRS2a, cell migration, and invasion. This indicates that binding of FGF9 to integrin is required for FGF9 signaling. Notably, R108E is a dominant-negative mutant, and effectively suppressed DNA synthesis, FRS2 phosphorylation and ERK1/2 activation induced by WT FGF9. This suggests that R108E has potential as a therapeutic in diseases in which FGF9 is involved. Previous studies showed that dominant-negative FGF1 mutant R50E bound to FGFR1 with an affinity comparable to that of WT FGF1 [26]. We predict that FGF9 induces integrin-FGF9-FGFR ternary complex on the cell surface, which is critical for FGF9 signaling. FGF9 binds to FGFR (high affinity receptor), and to integrins (low affinity receptor). It is highly likely that binding of FGF9 to FGFR is not sufficient for FGF9 signaling. It would be interesting to study if the R108E mutant of FGF9 suppress cancer proliferation by inhibiting FGF9 signaling (e.g., through FGFR3) in future studies.

Previous studies showed that several growth factors (e.g., FGF1, IGF1, fractalkine, and CD40L) require direct integrin binding and subsequent integrin-growth factor-cognate receptor ternary complex formation for signaling functions (ternary complex model) and growth factor mutants defective in integrin binding were defective in signaling functions, whereas they still bind to cognate receptors, and act as antagonists of growth factor signaling (growth factor decoys) [26, 28, 30-32]. We propose that integrins are common co-receptors of growth factors and the ternary complex model can be applied to many growth factor signaling. If this is the case, growth factor antagonists can be designed by screening growth factor mutants defective in integrin binding.

## Materials and Methods

DLD 1 human colorectal cancer and Colon26 mouse colon cancer cells were cultured in Dulbecco’s Modified Eagle Medium (DMEM)(Gibco, USA) containing 10% fetal bovine serum, 100 U/ml penicillin and 100 μg/ml streptomycin. NIH3T3 mouse embryonic fibroblasts, which were obtained from Bioresource Collection and Research Center, Taiwan, were cultured in DMEM containing 10% bovine serum (Gibco, USA) and MycoZap (LONZA, Switzerland) in an atmosphere of 95% air, 5% CO_2_. The antibodies were purchased from the following sources: rabbit anti-FRS2α (ProteinTech, USA), rabbit anti-phospho-FRS2α (Tyr-196) (Cell Signaling Technology, USA), rabbit anti-p44/42 MAPK (ERK1/2) (Cell Signaling Technology, USA), anti-phospho-p44/42 MAPK (p-ERK1/2) (Thr-202/Tyr204) (Cell Signaling Technology, USA), rabbit anti-FGFR1 (Cell Signaling Technology, USA), rabbit anti-FGFR2 (Sigma Aldrich, USA), and rabbit anti-FGFR3 (Novus Biologicals, USA).

### Synthesis of FGF9

The cDNA fragment encoding human FGF9 (LGEVGNYFGVQDAVPFGNVPVLPVDSPVLLSDHLGQSEAGGLPRGPAVTDLDHLKGI LRRRQLYCRTGFHLEIFPNGTIQGTRKDHSRFGILEFISIAVGLVSIRGVDSGLYLGMNE K GELYGSEKLTQECVFREQFEENWYNTYSSNLYKHVDTGRRYYVALNKDGTPREGTRTKRHQKFTHFLPRPVDPDKVPELYKDILSQS) was amplified by PCR using full length human FGF9 cDNA as a template and subcloned into BamH1/EcoR1 site of pET28aAmp vector, in which the kanamycin resistant gene was replaced with ampiciline resistant gene. Site-directed mutagenesis was performed using the QuickChange method [33]. The existence of FGF9 mutations was checked by DNA sequencing. The WT FGF9, R108E, K129E and K154E mutants were expressed in *E. coli* strain BL21(DE3) by isopropyl β-d-thiogalactoside (IPTG) induction and synthesized as insoluble proteins. The His-tag of proteins were used to purified proteins using Ni-NTA affinity chromatography in denaturing conditions (8 M urea). The Ni-NTA resin was washed with 0.5% Triton X-114 (Sigma, USA) to eliminate endotoxin before eluting the bound protein. The purified proteins were eluted in 250 mM imidazole/8 M urea. Purified proteins were diluted into refolding buffer (100 mM Tris-HCl, pH 8.0, 400 mM Arg, 2 mM EDTA, 0.5 mM oxidized glutathione, 5 mM reduced glutathione and PMSF) on ice. The dilution was kept for 16 hrs at 4°C with a slow stirring movement. Then the proteins were concentrated by ultrafiltration. Around 4 to 5 milligrams of purified proteins were obtained from one liter of bacterial culture. Purified proteins concentration was determined by measuring A280 and Bio-Rad protein Assay.

### Docking simulation

Docking simulation of interaction between FGF9 (PDB code 1IHK) and integrin αvβ3 (PDB code 1L5G) was performed using Autodock 3.05. In the present study, we used the headpiece (residues 1-438 of αv and residues 55-432 of β3) of αvβ3 (open-headpiece form, 1L5G.pdb). Cations were not present in αvβ3 during docking simulation [26, 34].

### Binding of soluble integrin αvβ3 to FGF9

ELISA-type binding assays were performed as previously described [30]. Briefly, wells of 96-well microtiter plates were coated with FGF9 by incubating for 30 min at room temperature and the remaining protein binding sites were blocked with BSA (heat-treated). Wells were incuvated with soluble αvβ3 (1 μg/ml) in Tyrode-HEPES buffer in 1 mM Mn^2+^ and incubated for 1 h at room temperature. After washing the wells with the sam buffer, bound αvβ3 was quantified using anti-β3 (mAb AV10) and HRP-conjugated anti-mouse IgG and peroxidase substrate.

### Cell migration assay

A polycarbonate filter of 8 μm pore size of the Chemotaxicell chamber (Kurabo, Osaka, Japan) was used to test cell migration. Lower side of the filter was coated with 10 μg/ml fibronectin (Asahi Glass, Tokyo, Japan) for 1h at room temperature. After washing, the chamber was placed into a 24-well cell culture plate, and the lower portion of the plate was filled with serum-free DMEM containing 50 ng/ml WT FGF9-or R108E FGF9. Cells were plated on the upper side of chamber and incubated at 37 °C for 6 h. Cells were fixed and visualized by crystal violet staining. The uncoated upper side of each filter was wiped with a cotton swab to remove cells that had not migrated through the filter.

Migrated cells were counted from the digital images of the stained cells, determining the mean number of cells counted per field. Results were expressed as means ± S.D. of the cell number.

### Invasion assay

Invasion assays were done in 8 μm pore size of the Chemotaxicell chamber coated with 100 μg/ml Growth factor reduced Matrigel (BD Biosciences, San Jose CA) for 3 h at 37°C and blocked with 0.1% bovine serum albumin in PBS. Cells were suspended in serum-free DMEM containing 0.1% BSA and plated in the upper chamber. The lower chamber was filled with DMEM containing 0.1% BSA and 50 ng/ml recombinant human WT FGF9 or R108E FGF9. Then, cells were allowed to migrate for 24 h. The top side of the filters was wiped with cotton swabs, fixed, and stained with 0.1% crystal violet. Images were taken by digital camera and counted using cell counting function of Image J software.

### BrdU Incorporation Assay

DNA synthesis or replication was monitored by measuring the Cell Proliferation ELISA BrdU kit (Roche). Cells were cultured on a Nunc 96-well plate (Thermo Scientific) at the density of 2×10^3^ cells/well. After 24 hrs, cells were rendered quiescent by incubation in serum free medium for 16 to 18 hrs and then stimulated with either WT FGF9 or FGF9 mutations for 16 hrs. BrdU labeling solution was added to each well concomitantly and those cells were incubated for 2 hrs in a CO_2_ incubator at 37°C in the presence of BrdU. Incorporated BrdU was detected using monoclonal mouse anti-BrdU antibodycojugated with HRP and tetramethyl-benzidine (TMB).

### Statistical Analysis

Data processing was performed using Prism 7 software (GraphPad, USA). All results were expressed as mean ± standard deviations. The statistical analysis of difference between two groups was analyzed by unpaired Student’s *t* test and difference between multiple groups was performed by one-way analysis of variance (ANOVA). P < 0.05 was considered to indicate statistically significant.

## Acknowledgements

This project was supported by pilot funding from the Comprehensive Cancer Center at UC Davis School of Medicine. This work is partly supported by the UC Davis Comprehensive Cancer Center Support Grant (CCSG) awarded by the National Cancer Institute (NCI P30CA093373) and the Pilot funding from the Department of Dermatology.

## Author Contributions

C-C Chang, YK Takada, Y Maekawa: data acquisition S. Mori, C-W Cheng and Y Takada: conceptualization, formal analysis, funding acquisition, and project administration.

## Conflict of Interest Statement

The authors declare that they have no conflict of interest.

**Supplemental Fig.**
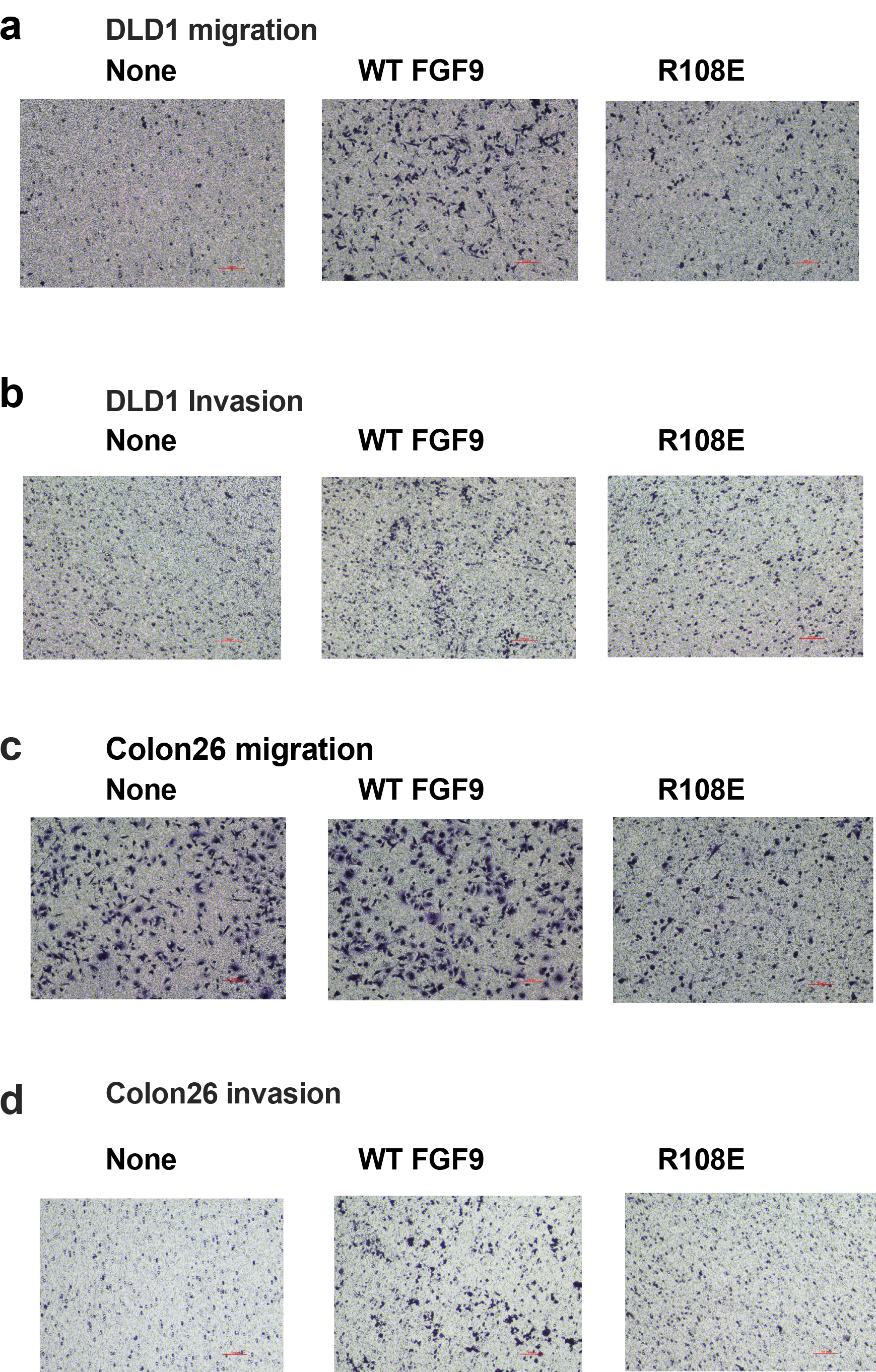

## Legend to supplemental figure

### Cell migration assay

A polycarbonate filter of 8 μm pore size of the Chemotaxicell chamber was used to test cell migration. Lower side of the filter was coated with 10 μg/ml fibronectin for 1h at room temperature. After washing, the chamber was placed into a 24-well cell culture plate, and the lower portion of the plate was filled with serum-free DMEM containing 50 ng/ml WT FGF9 or R108E FGF9. Cells were plated on the upper side of chamber and incubated at 37 °C for 6 h. Cells were fixed and visualized by crystal violet staining. The uncoated upper side of each filter was wiped with a cotton swab to remove cells that had not migrated through the filter. Migrated cells were counted from the digital images of the stained cells, determining the mean number of cells counted per field. Images were taken by digital camera and counted using cell counting function of Image J software.

### Invasion assay

Invasion assays were done in 8 μm pore size of the Chemotaxicell chamber coated with 100 μg/ml Growth factor reduced Matrigel for 3 h at 37°C and blocked with 0.1% bovine serum albumin in PBS. Cells were suspended in serum-free DMEM containing 0.1% BSA and plated in the upper chamber. The lower chamber was filled with DMEM containing 0.1% BSA and 50 ng/ml recombinant human WT FGF9 or R108E FGF9. Then, cells were allowed to migrate for 24 h. The top side of the filters was wiped with cotton swabs, fixed, and stained with 0.1% crystal violet. Images were taken by digital camera and counted using cell counting function of Image J software.

## Notes

### Competing Interest Statement

The authors have declared no competing interest.

